# Broccoli sprout Sulforaphane affected hemodynamics and aorta myogenic spontaneous rhythmic contraction in sanitized water uptake mice

**DOI:** 10.1101/2021.03.15.435382

**Authors:** Linmei Li, Lingfeng Gao, Ting Wang, Tong He, Jiaxing Li, Rifqah Kurnia Suwardi, Xuan Xu, Yang Wang

**Affiliations:** International nursing college, Hainan medical university, Haikou 571199, China; Extreme environment sports medicine laboratory, Hainan medical university, Haikou 571199, China; Bachelor of medicine and bachelor of surgery section, Hainan medical university, Haikou 571199, China

**Keywords:** Salinity water uptake, Broccoli sprout Sulforaphane, Carotid arterial pressure, Myogenic spontaneous contraction

## Abstract

Drinking seawater erodes water source will lead to hemodynamic changes in cardiovascular system. The erosion affected vascular biomechanics further interrupt the blood supply in arterial network. In this study, we investigated the carotid arterial hemodynamics in salinity water fed mice, and the relative spontaneous contraction of aorta preparation. The biological effect of Broccoli sprout Sulforaphane was assessed in intake hemodynamic changes. *Kunming* mice were randomly divided into seawater feeding group, seawater + Sulforaphane group, freshwater feeding group, fresh water + Sulforaphane group. After 4 weeks of feeding, the pressure waveforms of common carotid artery were analyzed *in vivo*. The enhanced common carotid arterial pressures were calculated according to the breakpoint of systolic pressure rising phase. The ejection time was calculated according to the dicrotic notch. *In vitro*, the isolated aorta biomechanical features were tested on a micro stepping platform. The passive tension and relative myogenic spontaneous contraction were evaluated. The results indicated that in salinity water fed mice heart rate, ejection period were significantly accelerated. The systolic pressure breakpoint of the ascending phase was significantly increased; however, the central aortic pressure augment index was decreased. *In vitro* study, the isolated aorta preparations indicated remarkable myogenic spontaneous contraction in salinity water fed mice. The spontaneous contraction indicated a significant cycle pattern, the waveform cluster changes regularly in one cycle, maximal amplitude of myogenic autonomic contraction increased significantly. Spontaneous contraction became more active, however cycle duration shortened. In biological effect of Broccoli sprout supplement, Sulforaphane was effective in reducing the heart rate, prolonging ejection period, improving systolic pressure and pulse pressure amplitude in salinity water fed mice. We concluded that long-term salinity water uptake can form a new hypertension model in mice, which can affect the changes of carotid artery hemodynamics and local blood supply. The Broccoli sprout Sulforaphane can improve the high systolic blood pressure and ejection period of artery, and its mechanism needs further study.

## Introduction

The problem of hypertension caused by long-term intake of salinity water has always been a concern of coastal countries and regions. Due to the melting of ice layer and the expansion of sea water caused by climate variation, the sea level average annual rises 3.2mm^[1]^, therefore cause high salinity erosion, which will have a lot of negative effects on domestic potable water sources. The high salinity water source is a prominent problem in coastal areas. High salinity drinking water can lead to cardiovascular disease, which is represented by hypertension and accompanied by increased vascular resistance with abnormal vascular function and structure. Although marine lipids, especially omega-3 polyunsaturated fatty acids (PUFAs), eicosapentaenoic acid (EPA) and docosahexaenoic acid (DHA) in infiltrate potable water, has a positive effect on preventing and alleviating chronic diseases. In recent years, it has been found that trace seawater can improve the balance of trace elements^[2]^, prolong the endurance of mice, reduce the oxygen consumption during exercise, and reduce the values of aspartate aminotransferase (AST), creatine kinase (CK) and creatine muscle enzyme isoenzyme (CK-MB) value has a certain effect^[3]^, improve the tolerance to high temperature^[4]^. However, the high incidence of hypertension caused by high salinity of seawater is still highly concerned, especially the abnormal lipid metabolism caused by abnormal copper and zinc metabolism, which is the cause of cardiovascular diseases such as hypertension, hyperlipidemia and atherosclerosis. Studies have shown that the blood copper, zinc, calcium and magnesium of rats fed with desalinated seawater were higher than those fed with fresh water, and the blood iron content decreased^[5]^. In the coastal areas of Bangladesh, drinking seawater erosion water cause maternal blood pressure increase, seriously affect maternal and fetal health. However, there was no significant correlation between the higher salt content water source living in inner Arizona and the rising prevalence of hypertension^[6]^, which suggested the different pathogenesis between hypertension caused by high salinity water eroded by seawater and hypertension caused by improper intake of daily edible salt, so the possible negative effects of multi-ionic components of seawater on cardiovascular system are considered. The studies shown that the average salt intake of 5g / day people had an increasing of systolic blood pressure 9mmHg who live in the coastal area of the bay of Bengal, while the systolic blood pressure caused by seawater salinity can reach to 120-139mmHg (even> 140mmHg in some cases, the diastolic blood pressure can reach to 80-89mmHg^[7]^. In the coastal areas of Vietnam, Bangladesh and India, salinization of drinking water has caused adverse effects on the health of more than 25 million people, resulting in the spread of hypertension and cardiovascular problem^[8]^. At present, scholars believe that millions of coastal residents are also at risk of hypertension and related diseases. It is urgent to conduct further research on seawater erosion and drinking water after desalination, so as to understand the health problems caused by marine water environment^[9]^.

The risk of high sodium intake in drinking erosion water is usually higher than that in food. However, the research in this area is basically blank, and the risk of seawater erosion of drinking water sources on blood pressure and cardiovascular disease is still unknown. Broccoli sprout Sulforaphane (SFN) is a molecule within the isothiocyanate (ITC) group of organosulfur compounds, which originally isolated from broccoli, a cruciferous vegetable. As the active component of its anti-inflammatory, SFN inhibit the proliferation and migration of hVSMCs induced by Ang II, and reduce the adhesion of monocytes to hVSMCs by reducing the levels of ICAM1 and VCAM1.

In order to provide a new experimental animal model for drug development, we designed the seawater water fed hypertension model for observing the hemodynamic characteristics of mice arterial system. We investigated the intake of SFN from cauliflower shoots, and explored the effect of Broccoli sprout crude extracted SFN in this hypertension model.

## Method

### Animal grouping

*Kunming* mice were provided and raised by the laboratory animal center of Hainan medical university (animal Certificate No.: 2018a044). 3-week-old male *Kunming* mice (n=20) were randomly divided into 1) Seawater fed and ordinary granulated food mice (hereinafter referred to as salinity group), which were further divided into salinity group (n = 5, sl) and salinity + supplement group (n = 5, sls). 2) The control group was fresh water and food feeding (hereinafter referred to as sanitary group), which were further divided into sanitary group (n = 5, sn) and sanitary + supplement group (n = 5, sns). In supplement feeding group, the Broccoli sprout crude extracted fluid gavage feeding twice per day and continued 4 weeks.

### Hemodynamic measurement

Mice anesthetized with 3% Pentobarbital Sodium (0.1ml/20g body weight, intraperitoneal injection), spray air to corneal reflexes with a pipette to confirm anesthetic effect. After fix the anesthetized mice on a wooden board in supine position, left common carotid artery was bluntly dissected through the midline incision of the neck. A polyethylene catheter (φ= 1 mm) was inserted into the common carotid artery from the incision centripetally. After ligation and fixation with silk thread, the blood pressure (PA) were monitoring through the common carotid artery. During monitoring, the pressure sensor should be consistent with the heart level. The characteristic waveforms of each stage of common carotid artery pressure are shown in **Figure 1**. The BL-420S data acquisition & analysis system recorded the pressure data. TM_wave software (ver 2.0) calculated central systolic pressure (CSP), central diastolic pressure (CDP), central pulse pressure (CPP) and carotid flow enhancement index (Faix (%) = cap / CPP * 100 (%)^[10]^. According to the obvious characteristics of the inflection point (PI) of the ascending branch of the common carotid artery systolic pressure at the end of inspiration, the inflection point of the common carotid artery systolic pressure at the end of inspiration in five breathing cycles was randomly selected to calculate the central augmentation pressure (CAP), and the ejection duration was calculated according to the distance between the rising starting point of systolic pressure and the notch of the beat wave.

**Figure 1.**
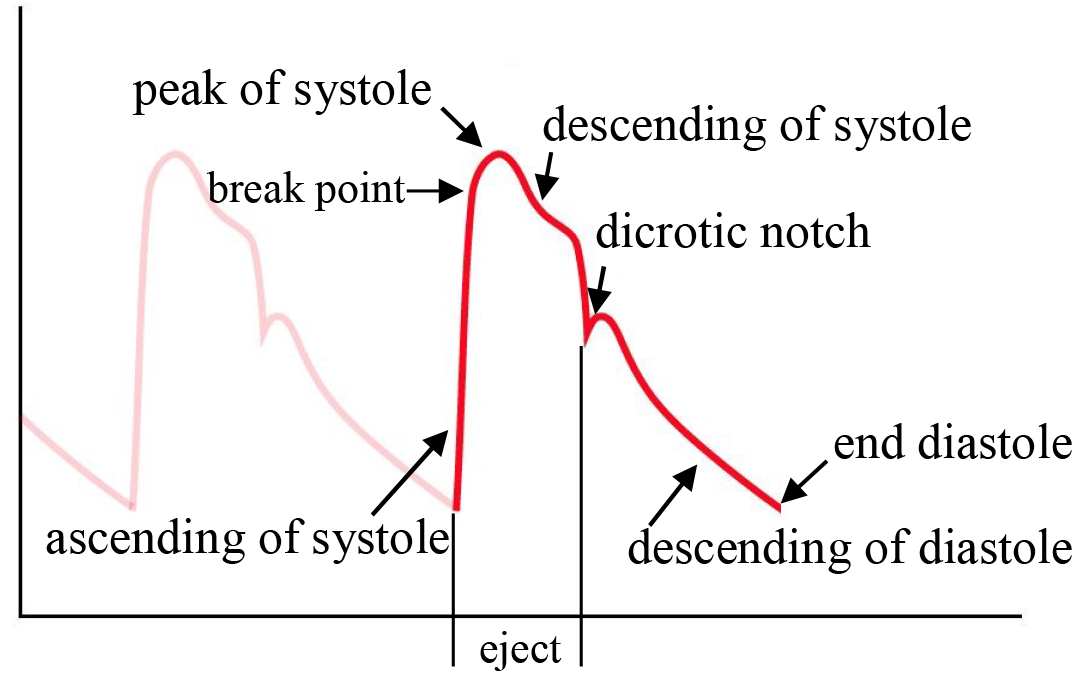
Carotid arterial waveform diagram

### Aorta mechanical lengthening tests in vitro

The proximal segments of aorta were isolated under the binocular dissecting microscope (pxs-2040, Shanghai Optical Instrument Factory, Shanghai, China). After removing the outer connective tissue, the aorta intima and smooth muscle layer were prepared. For further studies, the preparations were bath in *Ringer*’s solution with a constant temperature bath chamber (37 °C). Two preparations were prepared for each mouse.

The lengthening test was operated on a stepper motor driving roller screw platform. As shown in **Figure 2**, preparation side hooked on the glass probes. One side glass probe was fastened to a roller screw module; the other was connected to the reed of Wheatstone bridge-type piezoelectric strain sensor (Model number JH-2 10g, Beijing aerospace medical engineering institute, Beijing China). The roller screw module was installed on the vibration isolation platform (dst10-08, Jiangxi Liansheng experimental equipment) Equipment Co., Ltd., Shangrao, Jiangxi) to avoid environmental vibration interference. Preparations were kept horizontally on a 4°C chilled glass slide. The mechanical lengthening was operated by stepper motor which was controlled by Arduino Uno R3 board (Arduino, Allchips Ltd., Hong Kong). In order to obtain the steady mechanical lengthening in each stretch, the strengthen was determined by the rotating speed and the angles of stepper motor shaft which was driven by programmed pulse frequency of the Arduino board.

**Figure 2.**
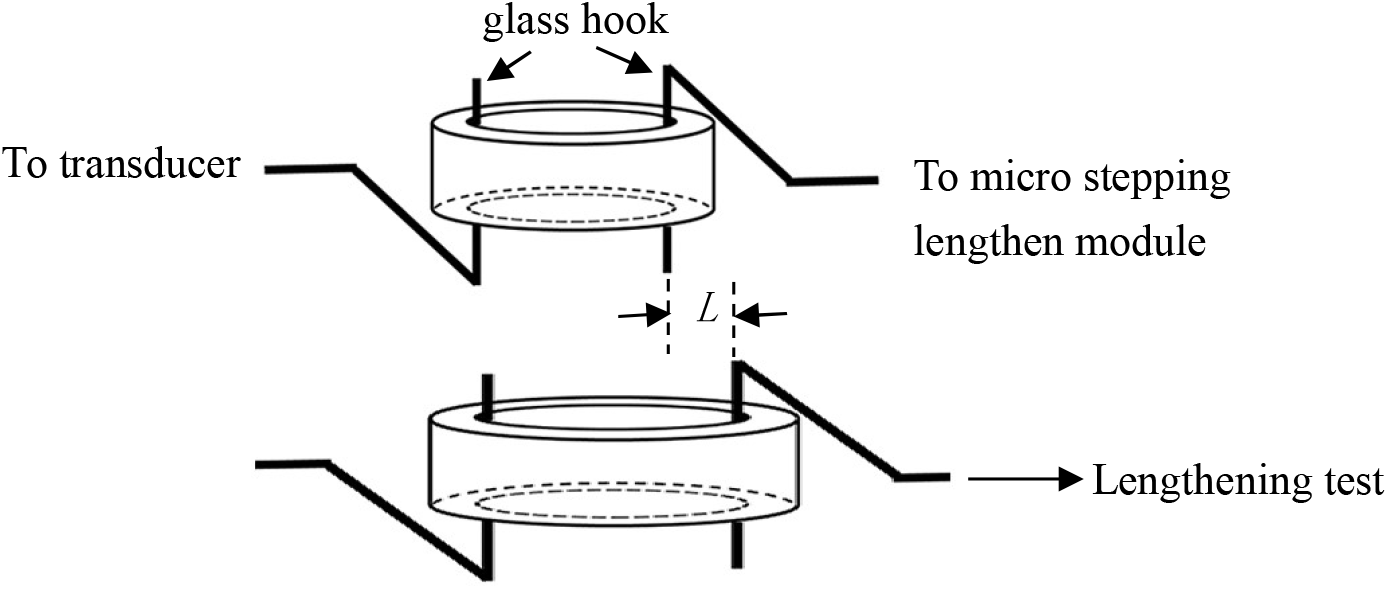
The schema of aorta lengthening loading test

As the procedure in **Figure 2**, the fixed preparations were slacked in *Ringer*’s solution, stabilize 5 min, then slowly manually turn the module, slightly stretch the preparation bearing 1g preload. The preparation length at this moment was defined as the initial length (*L_0_*). After the passive tension of the preparation was stabilized, the module was turned instantly. The preparations were passively elongated once based on its *L_0_* (**Figure 2**, the distance marked with *L* means the optimal length *L_0_* or rapid lengthening (*L_0_* + x). The passive tensions were traced and recorded by BL-420S data acquisition & analysis system. The myogenic spontaneous contraction and the characteristics of each contraction cycle was calculated and compared by TM_wave software (ver 2.0).

### Statistics

The data were presented as Mean ± Standard Error of Mean(−x ± SEM). Paired sample double population t-test was used for comparison between groups. Excel 2013 software (Microsoft Office Professional Plus 2013, Microsoft Corporation) for statistical analysis. *p* < 0.001 was statistically significant.

## Results

### The body weight between salinity and sanitary group

The average body weight was 34.70 ± 2.90g (n = 10) in salinity group, and 31.60 ± 1.90g (n = 10) in sanitary group. No significant difference between two group. After 4 weeks feeding, the body weight salinity groups significantly increased (53.77 ± 5.02g, n = 5, 155% increased; while 51.04 ± 4.72g, n=5, 162% increasing in sanitary group). However, in supplement groups, the body weight was not significantly increased (47.81 ± 3.59g, n=5, 138% increasing in salinity + supplement group; 49.32 ± 3.55g, n=5, 156% increasing in sanitary group). The body weight in supplement groups were significantly reduced (*p* < 0.001).

### The carotid arterial pressure waveform patterns in salinity mice group

In salinity group, the pressure waveform of common carotid artery changed significantly. The frequency of pressure waives significant increased. The pressure waives have a sharp peak of systole, that combining with a decrease rate of diastolic blood pressure accelerated. The ejection period (**ED**) shortened significantly. The amplitude of pulse pressure (**cPP**) increased significantly. Therefore, the break point of systole and the dicrotic notch were smoothing, and lost its normal pattern (**Figure 3**a sanitary mice, **Figure 3**b salinity mice). The characteristic wave pattern in salinity mice were the break point elevation significantly, resulting in the decrease of **cAP**. In salinity + supplement group, the elevations were rectified, which showed **cAP** increased. In sality mice, heart rate (HR) significant increased (742.99 ± 24.99 BPM and 698.12 ± 6.29 BPM in salinity group and salinity + supplement group, 448.36 ± 18.24 BPM and 427.19 ± 23.13 BPM in sanitary group and sanitary + supplement group).

**Figure 3.**
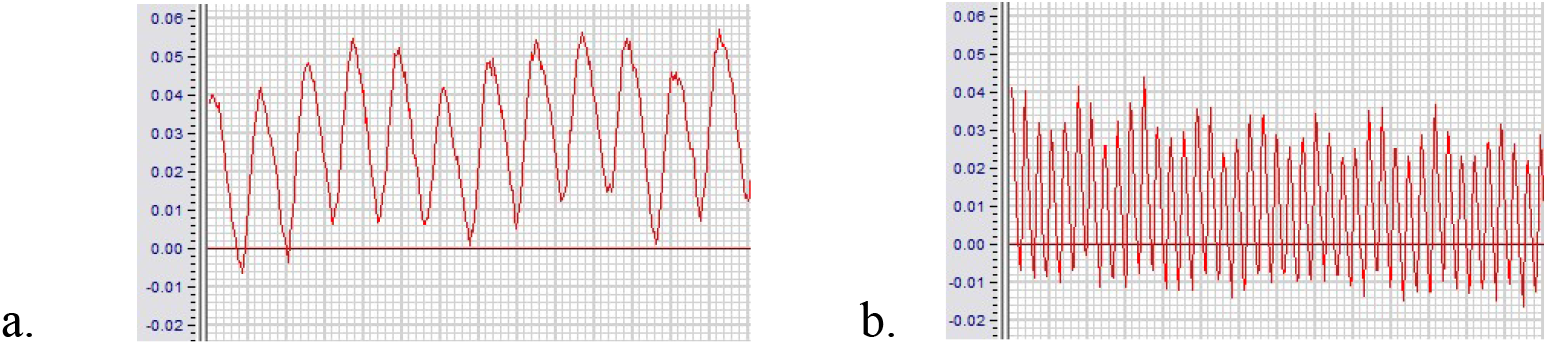
The carotid arterial pressure waveform in salinity and sanitary mice

### The cardiac output in salinity with supplement mice

The **CSP**, **CDP** and **cPP** in salinity group were significantly higher than those in sanitary group (21.42 ± 0.25kPa, 9.47 ± 0.22kPa and 11.95 ± 0.32kPa in salinity group; 18.27 ± 0.35kPa, 10.07 ± 0.23kPa and 8.20 ± 0.41kPa in salinity + supplement group). However, it was 13.13 ± 0.47 kPa, 6.52 ± 0.22 kPa and 6.61 ± 0.46 kPa, 11.53 ± 0.19 kPa, 6.43 ± 0.13 kPa and 5.10 ± 0.23 kPa in sanitary group and sanitary + supplement group.

**Figure 4** presented the statistic parameters comparison among the groups. The column bar with blank indicated the parameters in salinity and sanitary group, while the column bar with gray were the parameters in salinity + supplement and sanitary + supplement group. In **Figure 4**a, b and c, mark (a)*** was the statistic comparison result between salinity group and sanitary group, (b)*** was the comparison resupt between salinity + supplement group and sanitary + supplement group. The figures indicated the significant differences in **CSP**, **CDP** and c**PP** between salinity and sanitary mice (*** means *p* < 0.001); The mark (c) *** was the statistic comparison between salinity + supplement group and sanitary + supplement group (*** means significant difference, *p* < 0.001). The mark (d)*** in **Figure 4**c was the comparison between salinity group and sanitary group (*** means significant difference, *p* < 0.001), mark (e) *** was the ratio comparison between the salinity + supplement group and sanitary + supplement group (*** means significant difference, *p* < 0.001). Based on the above results, we concluded that salinity water feeding significantly increase **CSP** and **cPP** in mice circulation system. However, no statistic significant difference in **CDP**. SFN supplement reduce the **CSP**, improve the **cPP**, but no significant effect on **CDP**.

**Figure 4.**
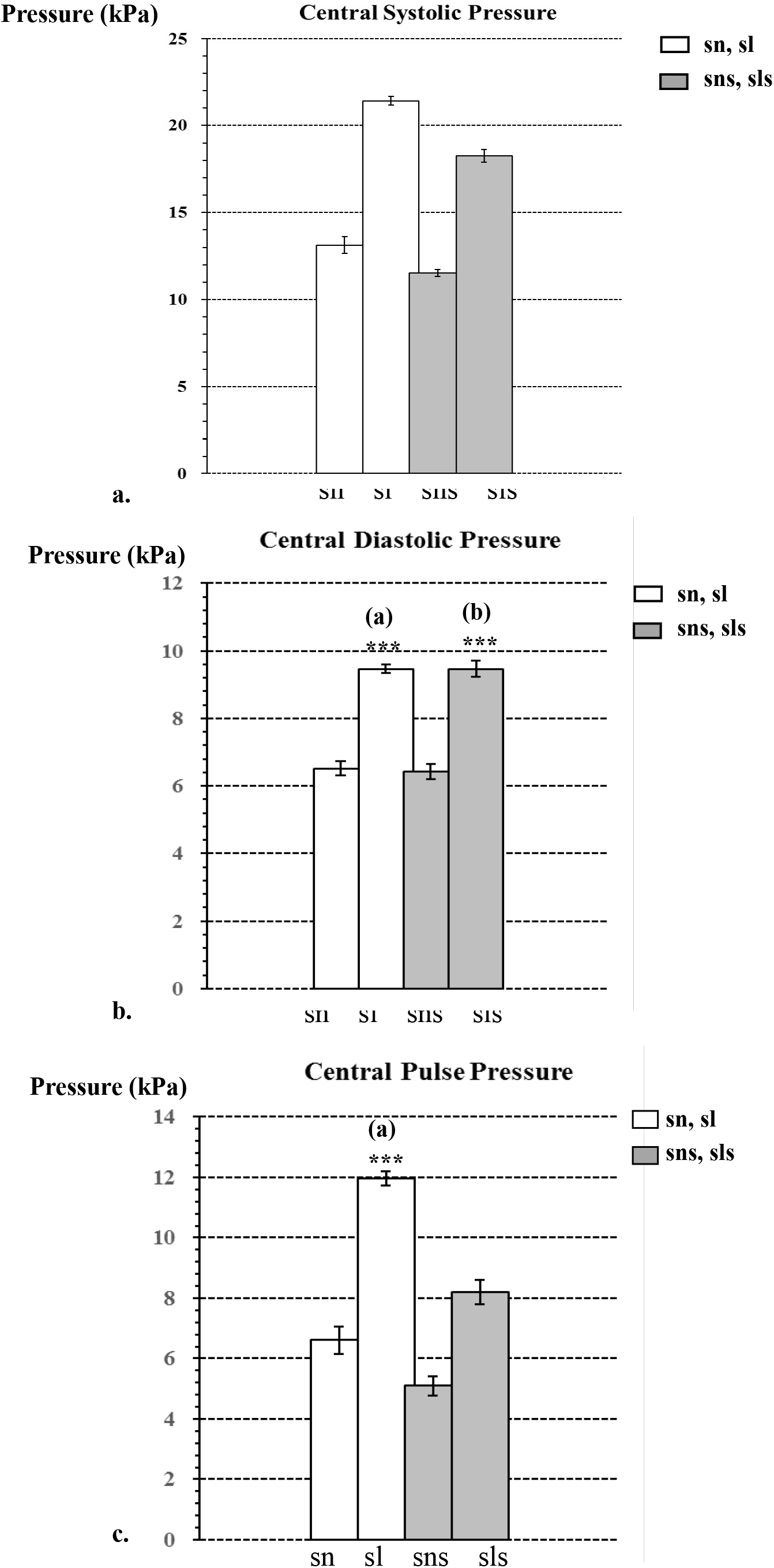
The comparison of carotid arterial **CSP**, **CDP** and **cPP** between salinity and sanitary groups

The **FAIx** were 71.82 ± 1.40, 89.73 ± 3.17 in salinity group and salinity + supplement group, and 80.24 ± 3.16, 91.02 ± 3.39 in sanitary group and sanitary + supplement group. **Figure 5** presented the comparison of **FAIx** between salinity and sanitary group, in which mark (a) *** was the statistic comparison result of **FAIx** between salinity group and sanitary group (*** means significant difference, *p* < 0.001); mark (b)*** and (c)*** were the statistic comparison result of **FAIx** between supplement and non-supplement in each group (*** means significant difference, *p* < 0.001). The results indicated that **FAIx** in salinity group was decreased, which may lead to the decrease of common carotid artery blood flow and local blood supply insufficiency in mice. However, supplement improved the **FAIx** in both group, which further suggest its improvements in common carotid artery blood flow and local blood supply insufficiency.

**Figure 5.**
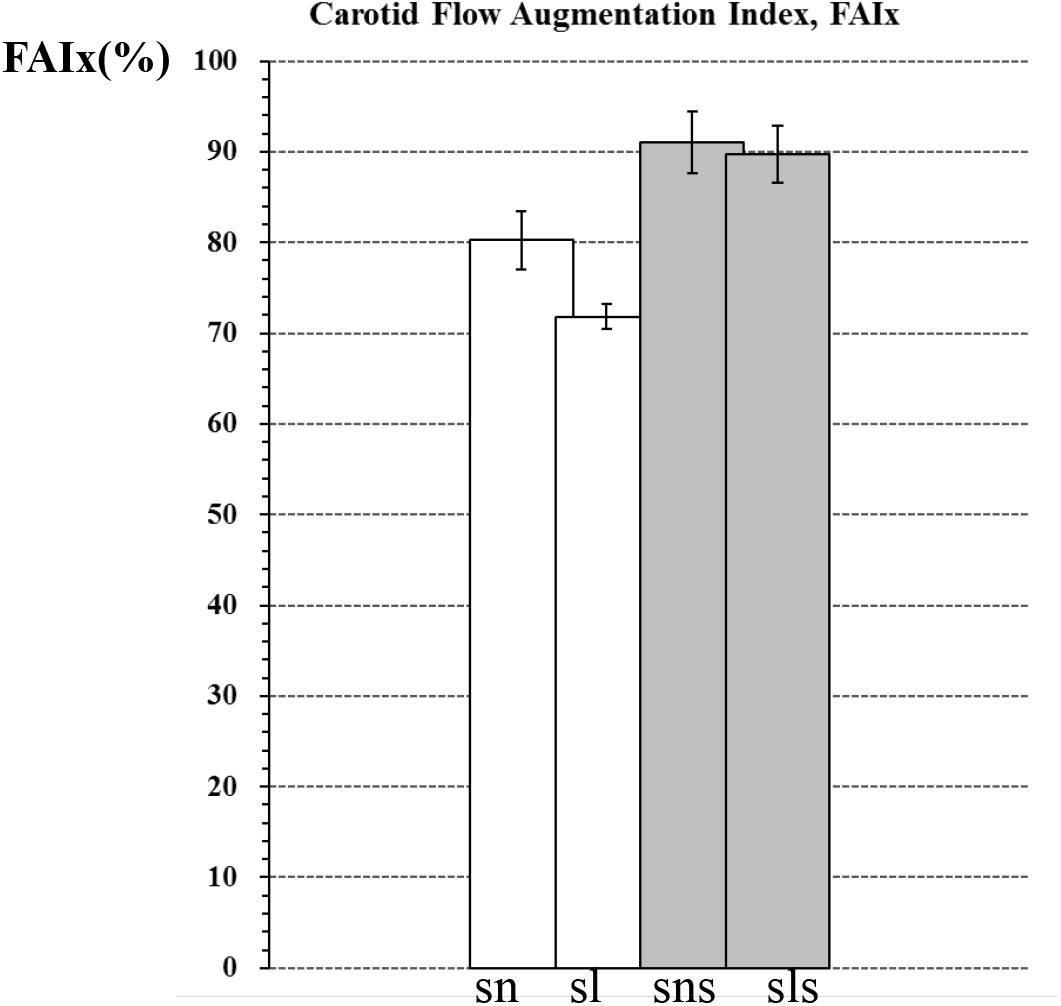
**FAIx** comparison in salinity and sanitary group

### The isolated carotid arterial preparation myogenic spontaneous contraction in salinity and supplement group

The maximal amplitude of myogenic spontaneous contractile was significantly increased in salinity mice, while the interval between the maximal amplitude was significantly shortened, and the frequency of wave clusters between the maximal amplitude was significantly increased (**Table 1**). This indicated seawater feeding evoked myogenic spontaneous contraction in aorta preparation. This myogenic spontaneous contraction response enhanced arterial resistance, leading to the decrease of **FAIx**, thereby reducing the blood flow and local blood supply. The effects of supplement was improvement of pulse pressure difference, but not the myogenic spontaneous contraction in salinity mice.

**Table 1.**
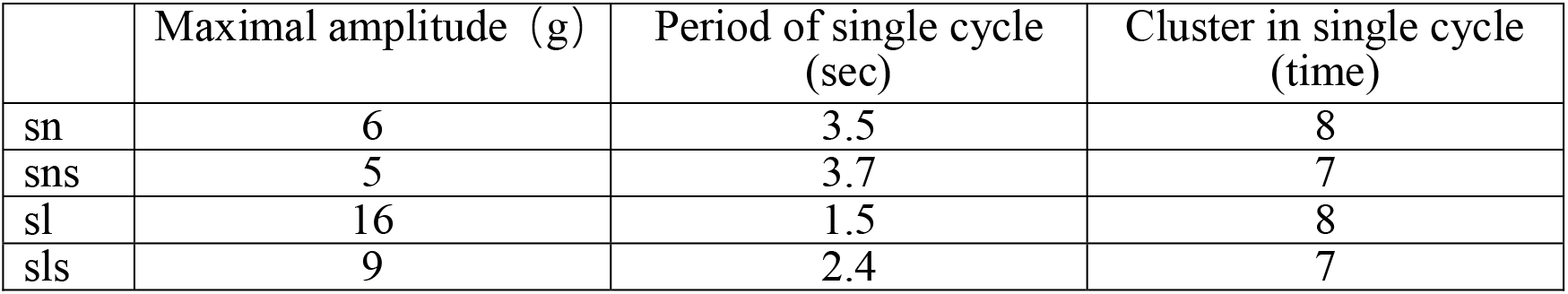
Aorta myogenic spontaneous contraction pattern in salinity and sanitary group

|  | Maximal amplitude (g) | Period of single cycle (sec) | Cluster in single cycle (time) |
| --- | --- | --- | --- |
| sn | 6 | 3.5 | 8 |
| sns | 5 | 3.7 | 7 |
| sl | 16 | 1.5 | 8 |
| sls | 9 | 2.4 | 7 |

## Discussion

Due to global climate variation, sea-level rise and various climate anomalies, salinization of water sources in coastal areas and its harm to the human circulation system have been paid more and more attention. Salinity in water is not expressed as a percentage in oceanography, but as a thousandth. The salinity of seawater is between 35 and 37 PPT, which can be quantified by measuring its conductivity. The higher the salinity is, the stronger the conductivity is. Salt comes from the dissolution of mineral water on the land and the deposition of solid and gaseous substances from the earth’s crust by rivers into the ocean, so the composition of electrolyte is more complex. Moreover, the change of salinity is related to the interaction of Ocean region, climate and global water cycle. In this experiment, the conductivity method was used to compare the conductivity between seawater and brine at room temperature, and then the salinity of seawater used in the experiment was calculated. The salinity of seawater used in the experiment was equivalent to 35% of brine (data is not shown).

It has been reported that Mg^2+^, Ca^2+^ and K^+^ in seawater are considered to have positive effects on the prevention of cardiovascular diseases. The effects of refining the mineral composition of deep seawater to 1000 hardness on the cardiovascular hemodynamics of rabbits after feeding showed that systolic blood pressure, diastolic blood pressure, pulse pressure, mean arterial pressure and total peripheral resistance decreased significantly. The slight increase of serum Mg^2+^ level in deep sea water group may not explain the inhibitory effect of mild hypertension^[11]^. However, the effect of normal seawater on cardiovascular system is negative. According to the research on the correlation between seawater and hypertension, it is believed that the high prevalence of hypertension in Alaska and Mekong Delta is due to the intake of seawater^[12, 13]^. The broad studies also show that there are genetic factors in the relationship between high salt diet and hypertension, and salt sensitive people are more likely to have hypertension. The salt sensitive hypertension mouse model is characterized by salt sensitive hypertension due to the decrease in water sodium retention and depletion of renal sodium excretion, and the decrease of salt excretion is related to the renin-angiotensin-aldosterone system (RAAS)^[14]^. In this study, we explicated the significantly different and unique carotid arterial blood pressure waveforms in the seawater feeding mice (salinity mice), which may belong to a new type of hypertension animal model. This included seawater feeding accompanied increasing of systole pressure and the significant myogenic spontaneous contraction pattern of aorta segment.

**FAIx** is a parameter describing the multiple relation between blood arteries and cardiac ventricles. It is related to the amplitude of the wave formed by the increase of left ventricular suction and the increase of blood pressure caused by the reflection of pressure wave. **FAIx** was more closely related to aortic pulse velocity, aortic compliance and elastic / muscle pulse velocity ratio. **FAIx** increased with the increase of the degree of arteriosclerosis. In this study, it indicated that **FAIx** was tightly relative to the myogenic spontaneous contraction in seawater feeding mice. The increased myogenic spontaneous contraction became more significant with the increase of arterial pressure and volume load, which may limit the blood flow in the arterial circulation. In recent scientific reports, it suggested that the changes of cerebral blood flow associated with large arteries may be related to **FAIx** variation^[15]^.

In seawater feeding mice, if the low **FAIx** relative to the cerebral blood supply insufficiency remains to need confirmed. Nevertheless, the phenomenon of cerebral blood supply insufficiency caused by low **FAIx** may provide a new research interests. Oral administration of seawater and gastrointestinal absorption of seawater are also effective ways to produce specific hemodynamic changes. Seawater can only be absorbed through the stomach and intestines to have a significant effect on the circulatory system, while the effect of other ways is very limited. According to the side effects of psoriasis patients treated with high salt seawater from the dead sea, 1142 psoriasis patients with hypertension were evaluated in the investigation. The decrease of diastolic and systolic blood pressure was not significant. The high salt environment in the dead sea had no significant side effects on the treatment of hypertension for psoriasis patients^[16, 17]^. Long term immersion in high salt seawater from the dead sea could improve blood pressure^[18, 19]^. Based on these reports, it can be concluded that oral uptake is the main way to cause seawater hypertension, while long-term seawater immersion and percutaneous contact will not have a negative impact on systemic blood pressure.

Sulforaphane is a metabolite of glucoraphanin (Grn), which in turn is the main glucosinolate (GLS) in broccoli. The production of sulforaphane is only possible when this is released due to plat injury. For example, chewing will chemically change the structure of glucoraphanin in conjunction with the enzyme myrosinase. Furthermore, this action will release glucose and sulfate, leaving the sulforaphane (SFN) molecule free to function. However, ITC and sulforaphane cytoprotective effect as an indirect antioxidant is associated with the fact that they can conjugate with glutathione (GSH), contributing to phase activation II enzymes and scavenging of ROS. Besides this, SFN can regulate the nuclear factor erythroid-derived 2-(NF-E2) related factor 2-(Nrf2-) antioxidant response element (ARE) pathway. Consequently, this action upregulates the expression of a range of antioxidant enzymes, including HO-1, NQO1, GST, γ-glutamyl cysteine ligase (GCL), and glutathione reductase (GR). The SFN-mediated protection against platelet aggregation has been well-documented. It is believed that SFN can decrease collagen-induced glycoprotein IIb/IIIa activation and thromboxane A2 formation. The ingestion of SFN resulted in a lower concentration of oxidized GSH, increased GR and GPx activity. Consequently, these improvements reflected in better endothelial relaxation and lower blood pressure.

In conclusion, we suggest that seawater feeding (sanitized water drinking) cause significant changes in the waveform of the common carotid artery, raise the break point of the ascending branch of systole pressure, reduce the central artery pressure, thus reducing the blood flow. As an important anti-inflammatory component, **br**occoli sprout sulforaphane reduced the systolic blood pressure of common carotid artery in seawater feeding mice. It could may be a potential compound to reverse the pathological changes of cardiovascular problem through its anti-oxidative stress pathways.

## Acknowledgement

This study was sponsored by national college student innovation and entrepreneurship project (No. 201911810021).

